# Laws of genome nucleotide composition

**DOI:** 10.1101/2023.09.09.557014

**Authors:** Zhang Zhang

## Abstract

Genome nucleotide composition is of fundamental significance in molecular evolution, genome organization and synthetic biology. Albeit studied for decades, it remains unclear whether there is any theoretical law underlying variable genome nucleotide composition across different species. From the mathematical viewpoint, here we propose three laws of genome nucleotide composition and validate their effectiveness on a large collection of empirical genome sequences across three domains of life. The three laws together provide a unifying framework that is able to unravel the mystery of genome-wide nucleotide composition variation across diverse species, paving the way towards a new era with quantitative insights for deciphering basic principles of life and further advancing theoretical biology.

**One-Sentence Summary:** The three laws of genome nucleotide composition provide a fundamental framework for molecular evolution, genome organization and synthetic biology.

## Introduction

Genome nucleotide composition, one of the most important characteristics at the genome-wide level, is usually expressed in terms of the proportions of four bases as well as their combinations in DNA molecule. It has been studied for decades (*1-3*) that genomes of different species are highly variable in their nucleotide composition (*4, 5*), as demonstrated that guanine-plus-cytosine content varies widely with a broader range from ∼20% to ∼80% (*6-9*). A body of empirical evidence has further accumulated that heterogeneity of genome-wide nucleotide composition in different species associates closely with a variety of factors, such as, genome size (*10*), phylogeny (*11, 12*), growth temperature (*13*), environment (*14*), origin of replication (*15*), transposable element (*16*), bacterial land colonization (*17*), codon/amino acid usage (*18-20*), cell fate control (*21*), and natural selection (*22-25*), etc. Although theoretical efforts have also been made in modeling the genome-wide nucleotide composition (*26-29*), it remains unclear whether there is any law underlying such variable genome nucleotide composition across different species. Theoretically, such law(s) would be desirable to be used as a fundamental framework for better understanding genome composition dynamics, molecular evolution, genome organization and synthetic biology. Built upon previous findings, here we propose three laws of genome nucleotide composition in a mathematical manner and demonstrate their effectiveness to formulate diverse genome nucleotide compositions in a large collection of 17,873 complete genome sequences across three domains of life.

### First law: the law of base pairing

The first law is Chargaff’s rules (*30-32*) that adenine (A) pairs with thymine (T) and guanine (G) pairs with cytosine (C), leading to *P*(A) = *P*(T) and *P*(G) = *P*(C) (Eqs. 1 and 2), where *P* is the proportion (probability) of any base as well as their combination. Such base pairing symmetry in any double-strand genome and each single strand corresponds to the Chargaff’s first parity rule and second parity rule, respectively. With biological, chemical and physical significances, the rules played a crucial role in the discovery of the double helix structure of DNA in 1953 (*33*) and laid profound foundations in advancing molecular biology and genomics (*34*). Despite the debate on the Chargaff’s second parity rule (*35, 36*), the first law holds valid as testified by nearly perfect linear regression in a wide diversity of genomes across the three domains of life (Fig. 1A and 1B).

**Fig. 1.**
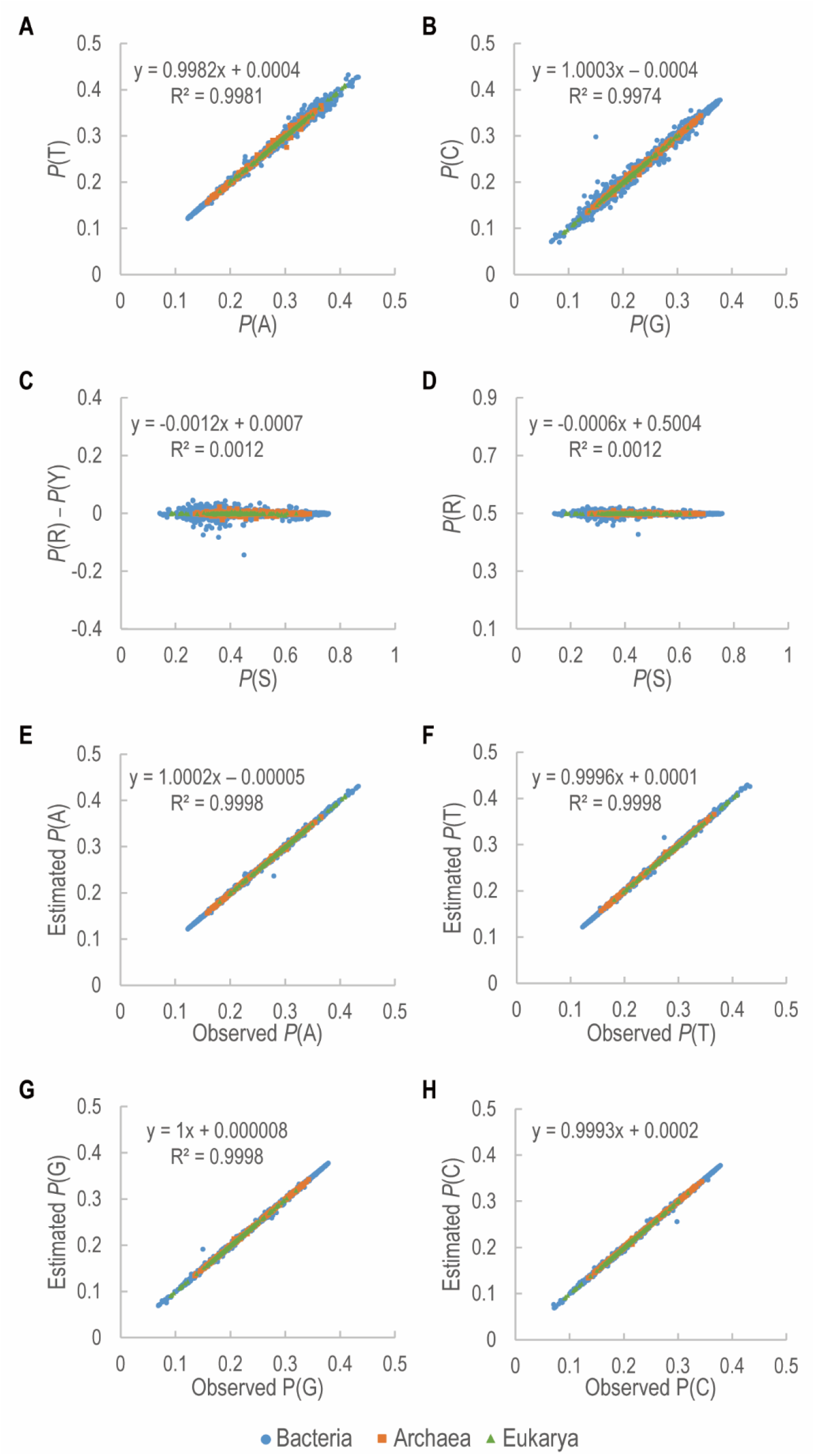
Genome-wide proportions of nucleotide composition across three domains of life. **(A)** Proportion of adenine (A) vs. proportion of thymine (T); **(B)** Proportion of guanine (G) vs proportion of cytosine (C); **(C)** Proportion of guanine-plus-cytosine content (S) vs. difference between proportion of purine content (R) and proportion of pyrimidine content (Y); **(D)** Proportion of S vs. proportion of R; **(E-H)** Observed vs. estimated proportions for A, T, G, and C, respectively. Observed proportions of the four bases were derived from a large collection of 17,873 complete genome sequences in NCBI RefSeq, including 313 in Archaea, 17,289 in Bacteria, and 271 in Eukarya. Estimated proportions of the four bases were quantified according to Eqs. 5-8. In all panels, each point represents the composition for a specific genome. All detailed information were summarized in Supplementary Table S1.

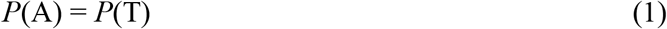

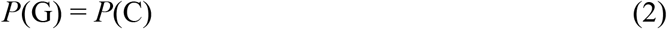

### Second law: the law of base equality

The second law states the base equality between purines (A and G) and pyrimidines (T and C). As a consequence of the base-pairing nature of DNA double helix, a 1:1 stoichiometric ratio of purines (R) and pyrimidines (Y) can be deduced. In other words, the proportion of R approximates the proportion of Y, namely, *P*(R) = *P*(Y). Additionally considering guanine-plus-cytosine (S or GC) and adenine-plus-thymine (W) contents, *P*(A) + *P*(G) + *P*(T) + *P*(C) = 100% can be further expressed as *P*(R) + *P*(Y) = 2*P*(R) = 2*P*(Y) = *P*(S) + *P*(W) = 100% (Eqs. 3 and 4).

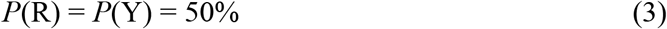

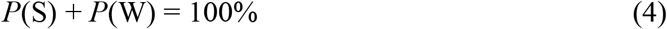

Conforming with the second law as well as previous findings (*4, 7, 8*), *P*(R) is observed to approximate *P*(Y) and fluctuate around 50% in diverse genomes belonging to Archaea, Bacteria and Eukarya (Fig. 1C and 1D). Meanwhile, considering that independence of two variables means that the occurrence of one variable does not affect the probability of the other, S, varying from ∼20% to ∼80%, is believed to be statistically independent from R, as indicated by linear regression slope very close to the optimum value of zero and intercept near 0.5 (Fig. 1D).

### Third law: the law of base composition

The third law expresses the principle of base composition. Suppose that the universal set of the four bases is Ω={A, T, G, C} and X and Y are two subsets of Ω, there are three common operations in set theory: (1) union of X and Y, viz., X∪Y, is the set of all elements that are a member of X or Y or both; (2) intersection of X and Y, viz., X∩Y, is the set of all elements that are members of both X and Y; (3) complement of X relative to Ω, viz., X^c^, is the set of all members that are not members of X. Thus, GC and purine contents can be denoted as S = G∪C and R = A∪G, respectively. Likewise, W = A∪T = S^c^ and Y = T∪C = R^c^. Because S and R form an independent pair as mentioned above, thus *P*(S∩R) can be quantitatively expressed as *P*(S) multiplied by *P*(R), namely, *P*(S∩R) = *P*(S) × *P*(R), which is also applicable to the other three pairs: S^c^ and R, S and R^c^, and S^c^ and R^c^ (for details see our previous study (*28*)). As a result, the proportion of each base can be quantitatively formulated as

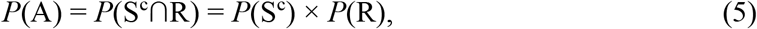

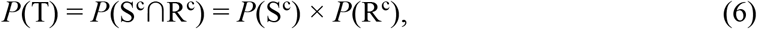

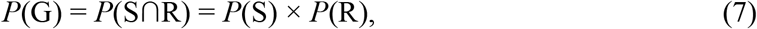

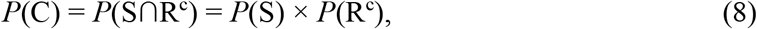

where *P*(S^c^) = *P*(W) = 1 – *P*(S) and *P*(R^c^) = *P*(Y) = 1 – *P*(R) according to Eqs. 3-4. Strikingly, if *P*(R) = 0.5, then *P*(A) = *P*(T) = *P*(S^c^)/2 and *P*(G) = *P*(C) = *P*(S)/2, which is a special case equivalent to the first law or Chargaff’s rules. Based on our empirical datasets, the third law is effective to quantify the base proportions very close to the observed ones, as signified by squared correlation coefficients very close to the optimum value of 1.0, linear regression slopes near the optimum value of 1.0 and intercepts approaching the optimum value of zero (Fig. 1E to 1H).

## Concluding thoughts

The first law of base pairing uncovers the complementary nature of DNA that is essential for its structure, stability and function, the second law of base equality reveals the equality relationship between purines and pyrimidines as well as the independence relationship between GC content and purine content, and the third law deduces the mathematical principle of each base composition that is of great utility for designing genomes with any specified composition. Together, the three laws provide a unifying framework for studying genome nucleotide composition (Fig. 2), which is not only able to unravel the mystery of various genome-wide nucleotide compositions across diverse organisms as validated in large-scale empirical genome sequences across the three domains of life, but also to detect organisms with unusual nucleotide composition that potentially experience complex evolutionary processes adapted to extreme environments (a case example is *Candidatus* Chazhemtobacterium aquaticus Ch65 (*37*) with strong disparity of 14.97% G and 29.83% C in Fig. 1B).

**Fig. 2.**
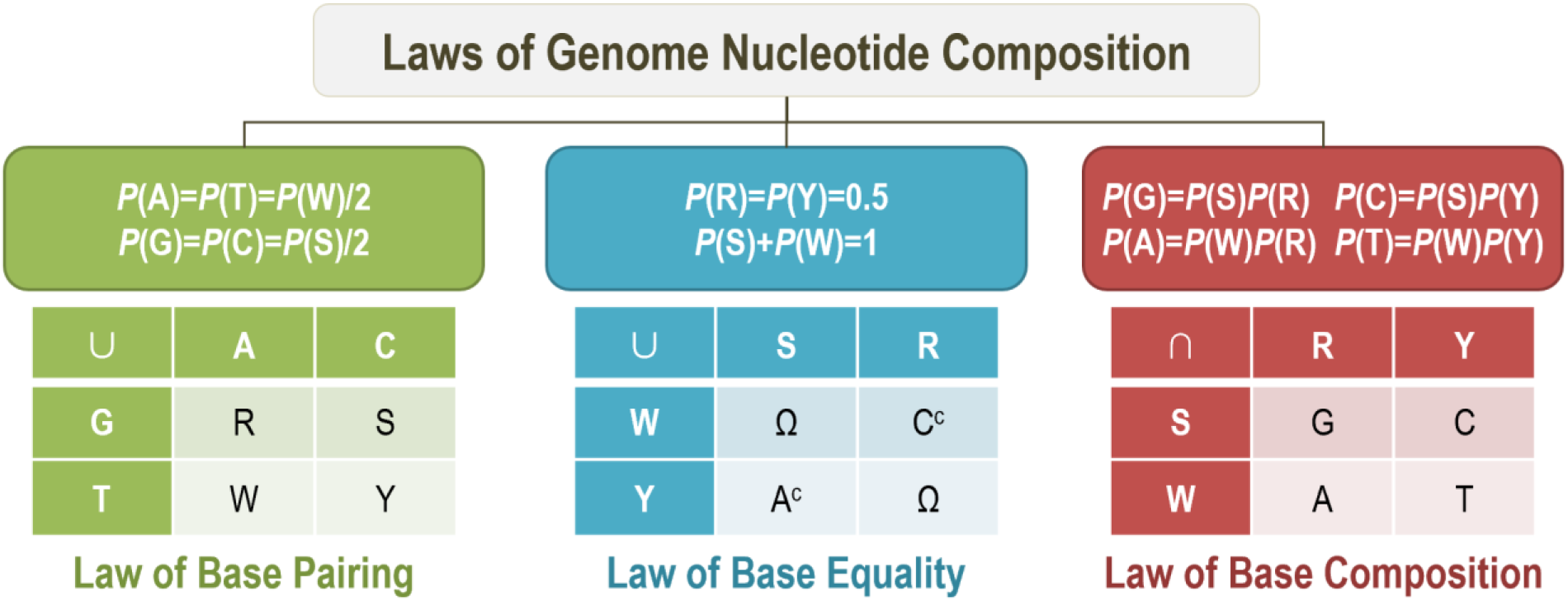
Schematic representation of laws of genome nucleotide composition. The laws are illustrated by equations and operations of intersection (∩), union (∪) and complement (^c^) on four bases—adenine (A), thymine (T), guanine (G) and cytosine (C), where *P* is the proportion (probability) of any base as well as their combination. The first law of base pairing states the complementary nature of DNA that leads to *P*(A)=*P*(T)=*P*(W)/2 and *P*(G)=*P*(C)=*P*(S)/2, where W=A∪T, S=G∪C, R=A∪G, and Y=T ∪ C. The second law of base equality reveals the quantitative relationships of *P*(R)=*P*(Y)=50% and *P*(S)+*P*(W)=100%, where R∪Y or S∪W is the universal set Ω={A, T, G, C}, R∪W={A, T, G}=C^c^ and Y∪S={T, G, C}=A^c^. The third law deduces the mathematical principle of each base composition given the independence relationship between S and R, viz., *P*(G)=*P*(S∩R)=*P*(S)*P*(R), *P*(A)=*P*(W∩R)=*P*(W)*P*(R), *P*(C)=*P*(S ∩Y)=*P*(S)*P*(Y), and *P*(T)=*P*(W∩Y)=*P*(W)*P*(Y), where W=S^c^ and Y=R^c^.

Nowadays (and in the foreseeable future), we are drowning in the deluge of multi-omics data growing at exponential rates, thirsting for fundamental theories to explain or predict a range of biological phenomena and to derive general laws from a number of empirical observations and experiments. From this perspective, the three laws, presented here as the union of theoretical and empirical work, potentially pave the way towards a new era with quantitative insights for studying genome organization and evolution, driving synthetic genome engineering and further advancing theoretical biology as a long-term objective on a par with theoretical physics.

## Authors’ contributions

ZZ conceptualized this idea, analyzed the data, plotted the figures, wrote the manuscript, acquired the funding support, and supervised the research. The author read and approved the final manuscript.

## Acknowledgments

I sincerely thank Haipeng Li, Yalong Guo, Yong Zhang, Jun Yu, Songnian Hu, Jingfa Xiao and Lina Ma for their valuable discussions on this work as well as Shuai Jiang and Zhao Li for their assistance in data collection. This work was supported by the National Natural Science Foundation of China (32030021) and International Partnership Program of the Chinese Academy of Sciences (153F11KYSB20160008).

## Supplementary information

**Table S1** Statistics of genome-wide nucleotide composition as well as other related meta information of 17,873 complete genome sequences across three domains of life.

